# Unmasking Epstein-Barr Role in Gastric Carcinogenesis: A Gene Expression Approach to Virus-Positive Tumors

**DOI:** 10.1101/2025.01.23.634441

**Authors:** Valéria Cristiane Santos da Silva, Diego Pereira, Daniel de Souza Avelar, Jéssica Manoelli Costa da Silva, Ronald Matheus da Silva Mourão, Rubem Ferreira da Silva, Davi Josué Marcon, Leandro Magalhães, Amanda Ferreira Vidal, Tatiane Neotti, Ana Karyssa Mendes Anaissi, Williams Fernandes Barra, Samir Mansour Casseb, Andrea Kely Campos Ribeiro dos Santos, Samia Demachki, Geraldo Ishak, Paulo Pimentel de Assumpção, Fabiano Cordeiro Moreira, Rommel Mario Rodriguez Burbano

## Abstract

Human gammaherpesvirus 4 or Epstein-Barr virus (EBV) is an oncogenic virus linked to malignancies like gastric adenocarcinoma. Notably, EBV infection induces genetic and epigenetic modifications that play a crucial role in oncogenesis and tumor progression, underscoring the importance of analyzing viral gene expression in the context of gastric cancer (GC) to elucidate its unique characteristics. This study aimed to perform a molecular characterization of EBV gene expression using next-generation sequencing (NGS). The analysis included human gene expression patterns in EBV-positive and EBV-negative samples and the expression of viral genes in EBV-positive samples. The study received approval from the Ethics and Research Committee of João de Barros Barreto University Hospital under reference number 47580121.9.0000.5634. It utilized 76 tumor tissue samples from patients with gastric cancer who had undergone surgical resection, and both fresh and paraffin-embedded samples were gathered for total RNA sequencing (RNA-seq) and in situ hybridization (ISH). The RNA-seq was conducted in a pair-end manner on the NextSeq® platform (Illumina®, US). The NextSeq® 500 MID Output V2 kit - 150 cycles (Illumina®) were utilized following the manufacturer’s instructions. ISH targeting RNA-1 of EBER1 (Y5200, DAKO, Carpinteria) was performed using the automated Dako system. Molecular characterization was conducted using the Kraken2 software. Subsequently, to elucidate the mechanisms through which EBV may influence gastric cancer, we analyzed the patterns of human gene expression in EBV-positive and EBV-negative samples. Of the 76 samples, 8 were classified as EBV-positive according to the applied methodology. Our analysis identified approximately 834 differentially expressed genes, 92 of which exhibited an AUC > 0.85. These genes are implicated in tumor progression, cellular metabolism, and both innate and adaptive immune responses. Additionally, viral genes expressed in the positive samples were evaluated, and we found manifestations of both lytic phase and latent phase genes. Finally, our study presents an efficient strategy for molecular classification of EBV-positive gastric cancer based on NGS and shows the effects of EBV on human gene expression.

**Author summary:** In our study, we explored how EBV influences the development of stomach cancer. EBV is a virus known to be linked to various cancers, including gastric cancer, and it can alter the behavior of both human and viral genes within infected cells. To investigate this, we analyzed tissue samples from 76 patients with stomach cancer, focusing on differences between samples with and without EBV. Using advanced sequencing technology, we identified over 800 genes that behave differently in EBV-positive cancers. These genes are involved in critical processes like how cells grow, how the immune system responds, and how energy is produced within cells.

We also examined which EBV genes were active in the cancer samples and found evidence of both dormant and active phases of the virus. Our work demonstrated how EBV may contribute to stomach cancer and suggests new ways to classify and understand this disease. By uncovering these details, we hope to pave the way for more targeted treatments in the future.

## Introduction

Gastric cancer (GC) ranks fifth in global cancer incidence and fourth in cancer-related mortality[1]. In Brazil, the most prevalent form of gastric cancer is gastric adenocarcinoma (GA), accounting for 95% of cases, with the highest incidence rates observed in the northern and northeastern regions of the country[2]. The primary causes of GC are linked to genomic regulatory failure[3], environmental factors such as dietary habits, excessive alcohol and tobacco consumption, and infections by microorganisms, including *Helicobacter pylori* (*H. pylori*) and Epstein-Barr virus (EBV)[4]. GC is classified according to histological and molecular characteristics to enhance understanding and treatment approaches. The Cancer Genome Atlas (TCGA) proposed a molecular classification in 2014, which categorizes GC into four subtypes: microsatellite instability (MSI), genomically stable (GS), chromosomal instability (CIN), and EBV-positive (EBV+) tumors[5].

Epstein-Barr virus (Human gammaherpesvirus 4), a member of the *Herpesviridae* family, is the first virus identified to have oncogenic properties associated with malignancies such as nasopharyngeal carcinoma, Hodgkin lymphoma, Burkitt lymphoma, and gastric cancer[6]. Notably, EBV infection induces epigenetic modifications that play a crucial role in oncogenesis and tumor progression, underscoring the importance of analyzing viral gene expression in the context of gastric adenocarcinoma to elucidate its unique characteristics[7]. EBV infection primarily occurs through the oral route, predominantly affecting B lymphocytes and epithelial cells. The virus initially replicates in the epithelial cells of the oropharynx, facilitating its spread to B cells, a process enhanced by the interaction between the viral glycoprotein gp350 and the *CD21* receptor[8].

EBV establishes a latent infection, utilizing proteins such as *EBNA-1* to evade immune detection[8]. The virus alternates between lytic and latent phases; the lytic cycle involves viral genome replication and the production of new virions, regulated by immediate-early, early, and late genes[9]. Latency is characterized by distinct gene expression patterns associated with various disease states[10]. Type III latency is initially established in infected B cells, but T cells typically recognize and eliminate these cells. EBV can also establish type 0 latency in memory B cells, characterized by minimal gene expression[8]. In gastric cancer, EBV is often associated with an intermediate latency phase between types I and II, characterized by limited protein expression and low-intensity epigenetic modifications[11]. The consequences of EBV infection are significant, as the virus exhibits both oncogenic and immunogenic activities[11,12]. The virus alternates between lytic and latent phases; the lytic cycle involves viral genome replication and the production of new virions, regulated by immediate-early, early, and late genes[9]. Latency is characterized by distinct gene expression patterns associated with various disease states[10]. Type III latency is initially established in infected B cells, but T cells typically recognize and eliminate these cells. EBV can also establish type 0 latency in memory B cells, characterized by minimal gene expression[8]. In gastric cancer, EBV is often associated with an intermediate latency phase between types I and II, characterized by limited protein expression and low-intensity epigenetic modifications[11]. The consequences of EBV infection are significant, as the virus exhibits both oncogenic and immunogenic activities[11,12].

A significant feature of Epstein-Barr virus-associated gastric cancer (EBVaGC) is DNA hypermethylation in gene promoter regions, affecting tumor suppression and processes like cell cycle regulation, cell adhesion, metastasis, apoptosis, and DNA repair. This hypermethylation occurs in both the host and viral genomes and is linked to carcinogenesis[13]. Aging, smoking, and viral proteins like *EBNA1* and *LMP2A* can trigger hypermethylation through DNA methyltransferase overexpression, potentially serving as a host defense mechanism against EBV infection[6,14]. Expressing latent cycle proteins such as *EBERs, EBNA1,* and *BARTs* characterizes EBVaGC. *EBERs* are involved in cell proliferation, resistance to apoptosis, and miRNA expression modification, particularly in suppressing the *CDH1* gene[6].

*EBNA1*, crucial for viral replication and epigenetic deregulation of tumor suppressor genes like gastrokines, also exhibits oncogenic potential, though its expression in gastric cancer is often reduced[14]. *BART* transcripts, including *BARF1*, are linked to chemotherapy resistance and the silencing of proliferation-related genes. *BARF1* promotes gastric cancer cell proliferation through the NF-kB/cyclin D1 signaling pathway[6,14]. *BARTs* also regulate *LMP1* and *LMP2A*, impacting viral latency and reactivation, with *LMP1* influencing proliferation, cell migration, and epigenetic changes in the host genome.

Examining viral gene expression in tumor cells is essential to determine the predominant stage of the viral cycle in tumor samples and its impact on cancer development. This study aims to elucidate the characteristics of EBVaGC, as identifying EBV presence in GC samples and analyzing its role in carcinogenesis hold substantial clinical relevance. Patients with a confirmed EBV infection exhibit distinct molecular and histological features, often necessitating tailored therapeutic approaches.

## Results

### ISH and molecular classification of EBV

For the molecular classification of EBVaGC, the criterion established to consider a sample positive was the detection of viral gene reads ≥ 100. The results obtained by the RNA-Seq method were compared with the *in situ* hybridization data performed by the pathology department of the João de Barros Barreto University Hospital to assess the parity of the results (**Fig 1**). Of the 76 selected tumor samples, 8 samples were considered positive for the presence of the virus by both methods. Among the samples tested by the ISH test, 1 sample did not demonstrate sufficient reads to be considered positive by the RNA-Seq method. The other samples considered negative by ISH showed viral gene reads ≤ 100.

**Fig 1.**
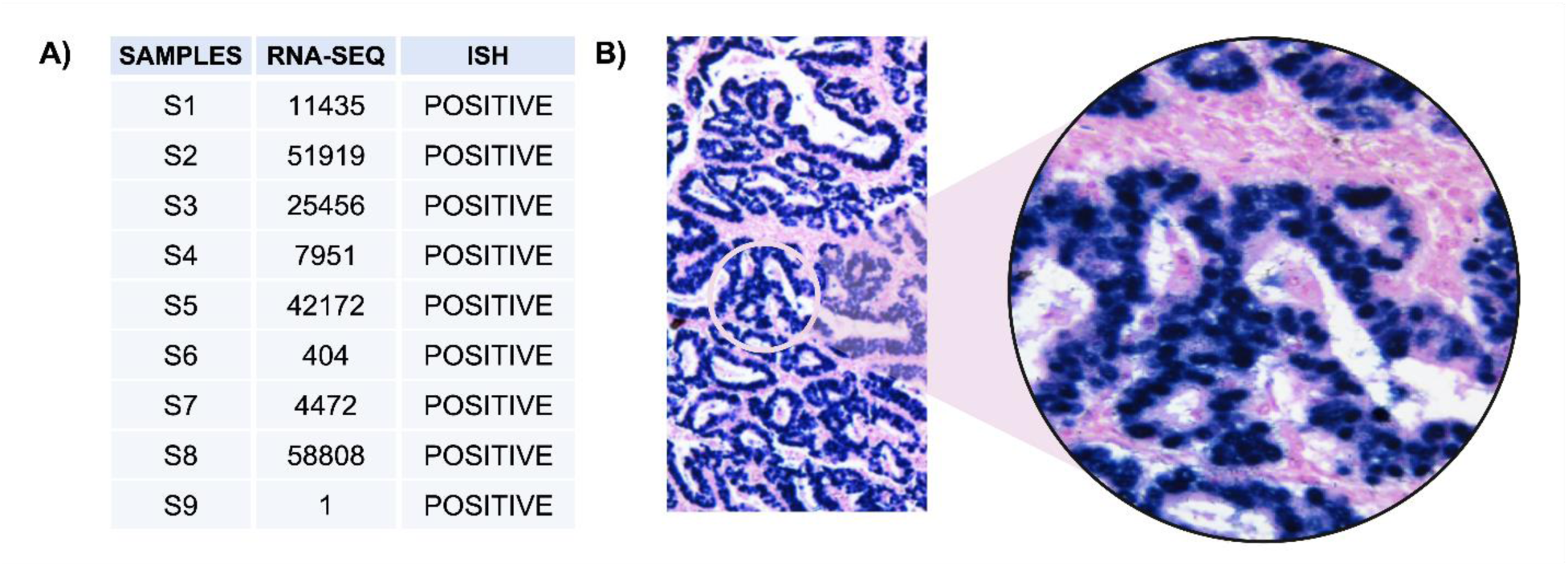
In situ hybridization of EBVaGC samples. (A) EBV-positive samples by ISH and number of EBV reads by RNA-Seq. (B) Histological slide representing a sample positive for EBV by ISH.

### Analysis of Differential Gene Expression and functional enrichment analysis of DE genes

After classifying the samples, a differential gene expression profile analysis was conducted between EBVaGC and EBVnGC patients. In this analysis, 834 human genes were found to be differentially expressed between the samples based on the established criteria for differential expression (DE) genes. Among these genes, 612 were downregulated, while 222 were upregulated. To evaluate which DE genes have higher specificity and sensitivity in distinguishing between EBVaGC and EBVnGC samples, we selected genes with an Area Under the Curve (AUC) value > 0.85. Of the 834 DE genes, 93 were selected based on the AUC cutoff value, and among them, the top 20 with the highest AUC values were investigated for their influence on carcinogenesis. The genes *GBP5* and *TYMP* were highlighted as they exhibited an AUC > 0.95 **(Fig 2)**.

**Fig 2.**
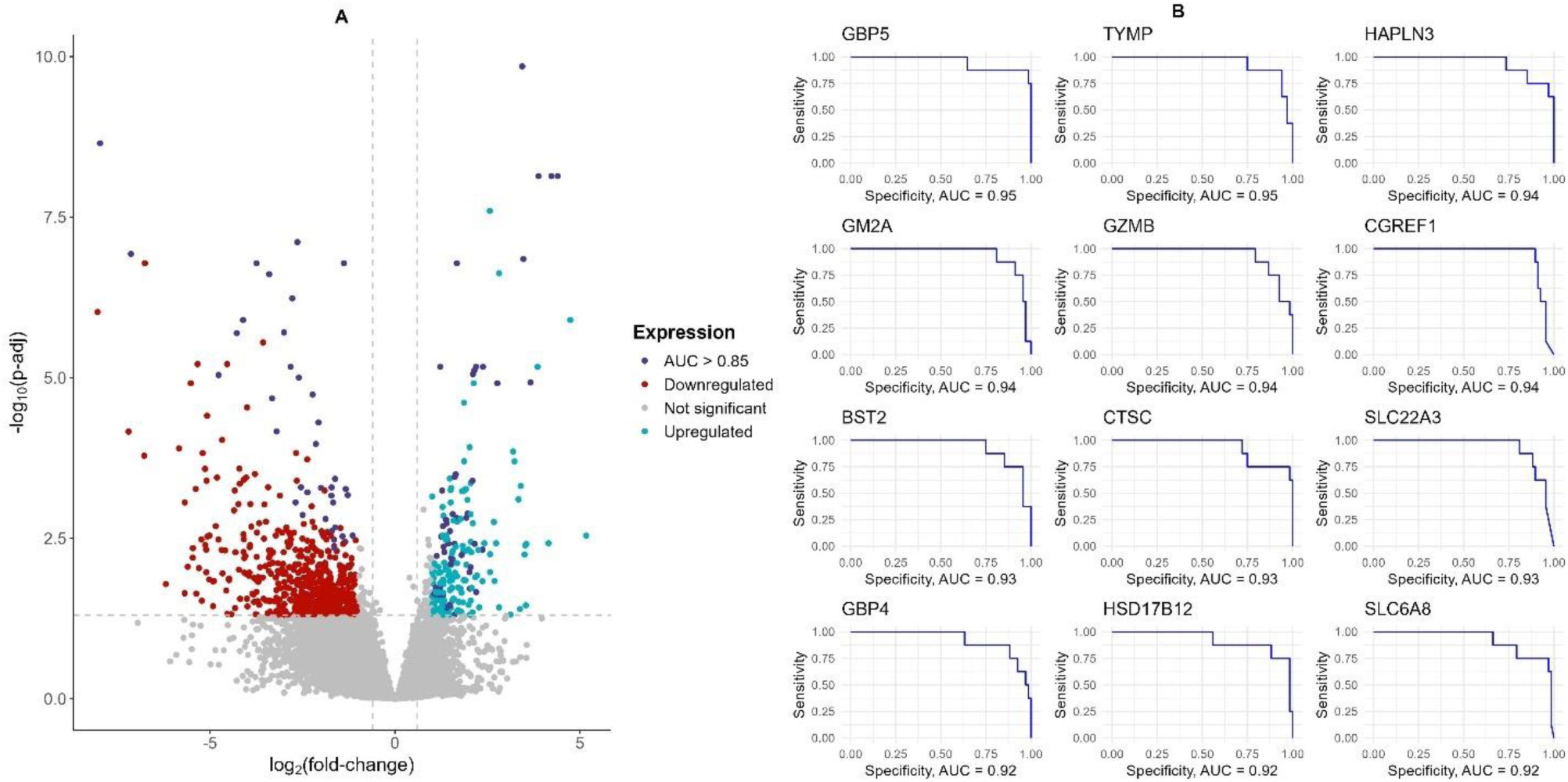
Differentially expressed genes in EBVaGC. (A) Volcano plot showing DE genes between EBV-positive and EBV-negative samples. (B) ROC curve plots representing the AUC of the top 12 DE genes.

To better visualize the gene expression patterns between EBVaGC and EBVnCG samples, we selected only the genes with AUC > 0.85 and conducted an unsupervised hierarchical clustering analysis (**Fig 3**). We observed that EBVaGC samples exhibit a distinct gene expression pattern compared to EBVnGC samples, as evidenced by the formation of clusters. This indicates that certain groups of genes show negative regulation in EBVaGC samples compared to EBVnGC samples. Additionally, the heatmap demonstrates that the samples were grouped based on similarities in their expression patterns.

**Fig 3.**
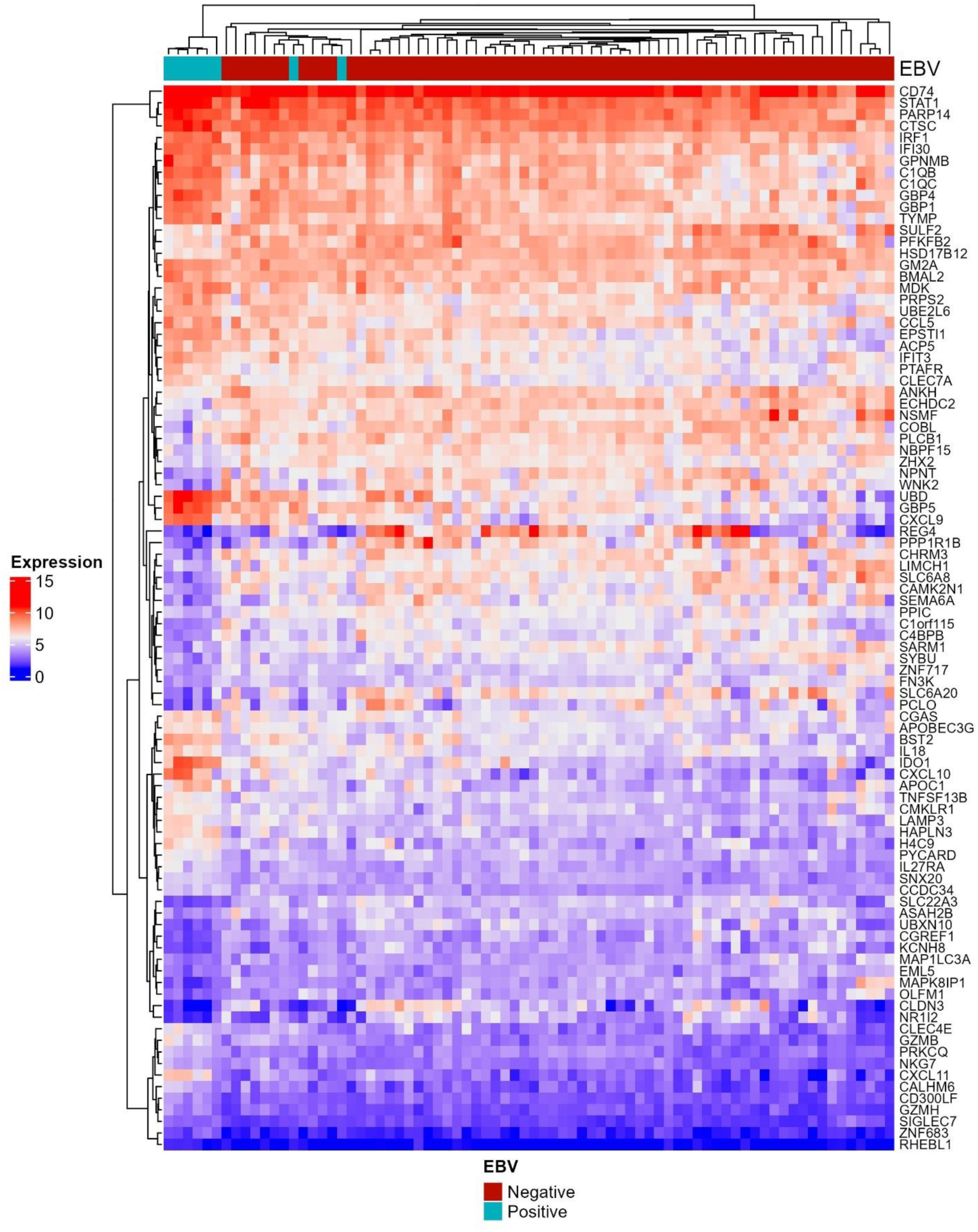
Heatmap of the expression patterns of DE genes with AUC > 0.85.

To better understand the functions and influences of DE genes, we performed ontology analysis using Gene Ontology (GO) with the ClusterProfiler package, which yielded significant terms related to biological processes. Among the primary enriched terms, several related to immune response were highlighted, including T-cell activation regulation, positive regulation of defense response, response to the virus, T-cell proliferation regulation, and lymphocyte proliferation regulation (**Fig 4**).

**Fig 4.**
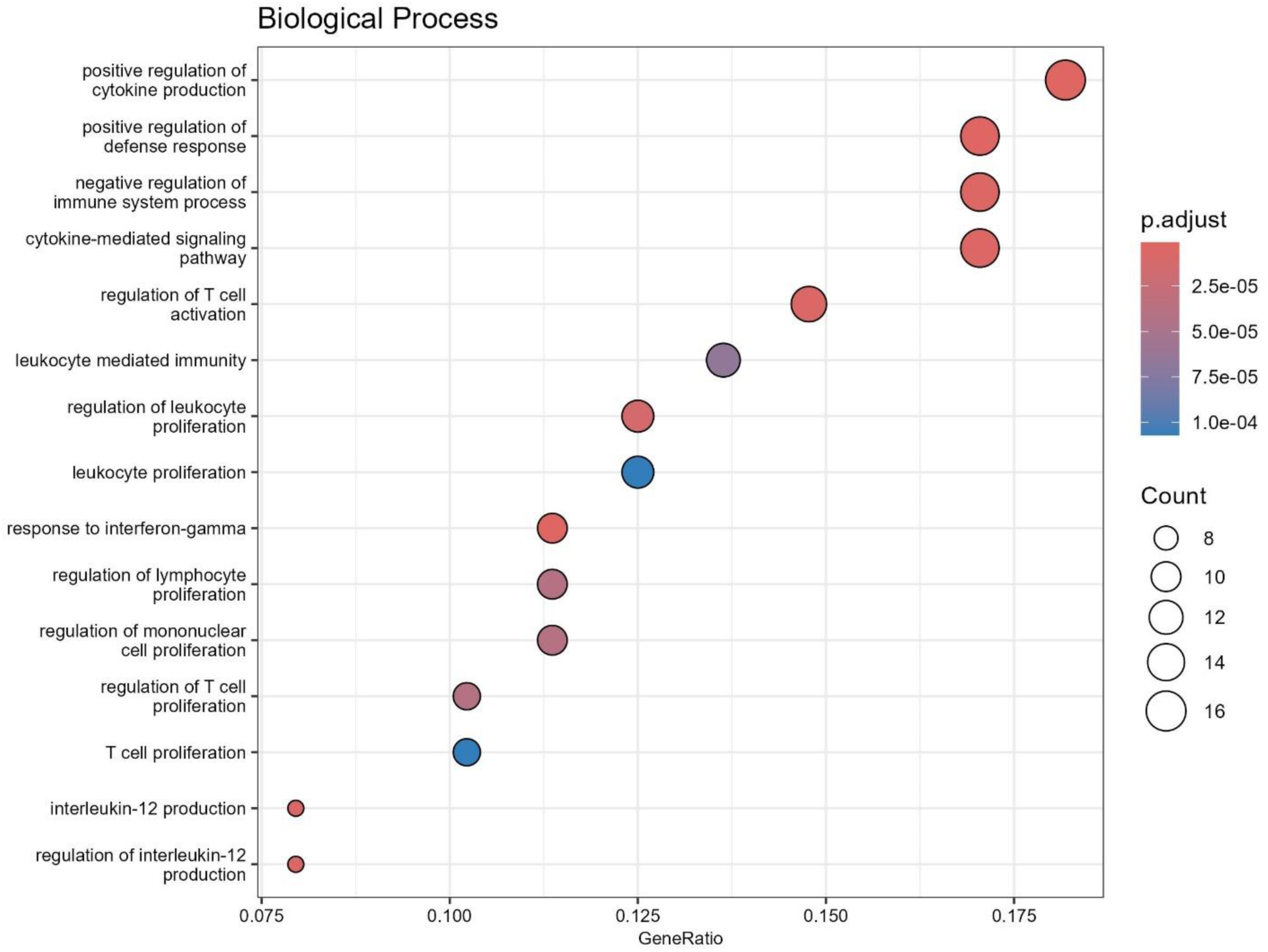
Gene ontology of biological processes enriched for DE genes.

### Analysis of Viral Gene Expression

Our analyses focused on evaluating the expression of viral genes in EBV-positive samples. Among the most frequently expressed genes, we identified *RPMS1, A73*, and *LMP-2B* as present in all samples. We also observed high expression of *EBER-1 and EBER-2*, which are important non-coding RNAs for virus identification in ISH tests (**Fig 5**). Subsequently, we evaluated the viral gene expressions by sample. Sample S3 has the highest expression of EBV genes, while sample S1 has an expression of only 4 viral genes (**Fig 6**).

**Fig 5.**
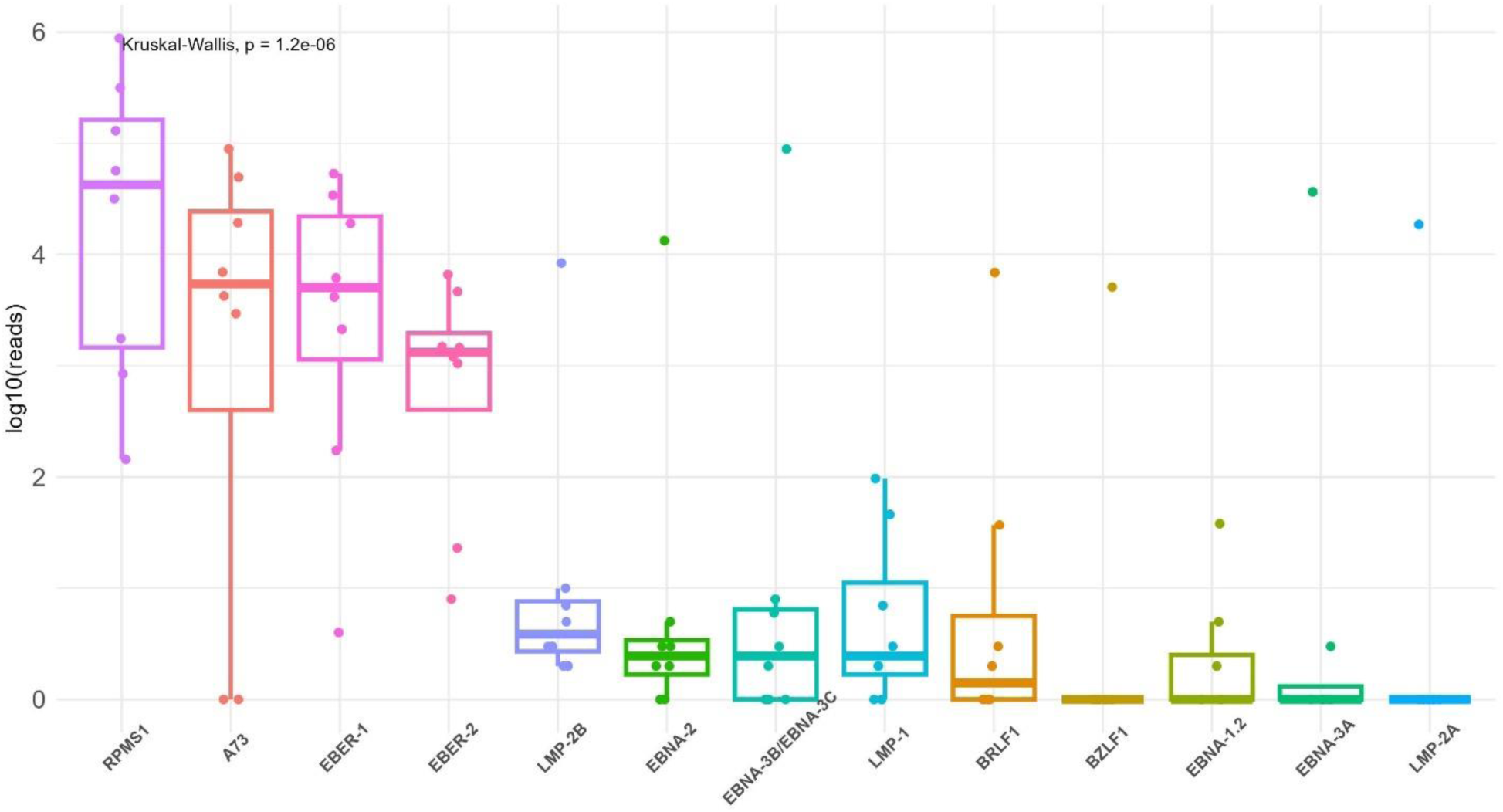
Expression of EBV viral genes.

**Fig 6.**
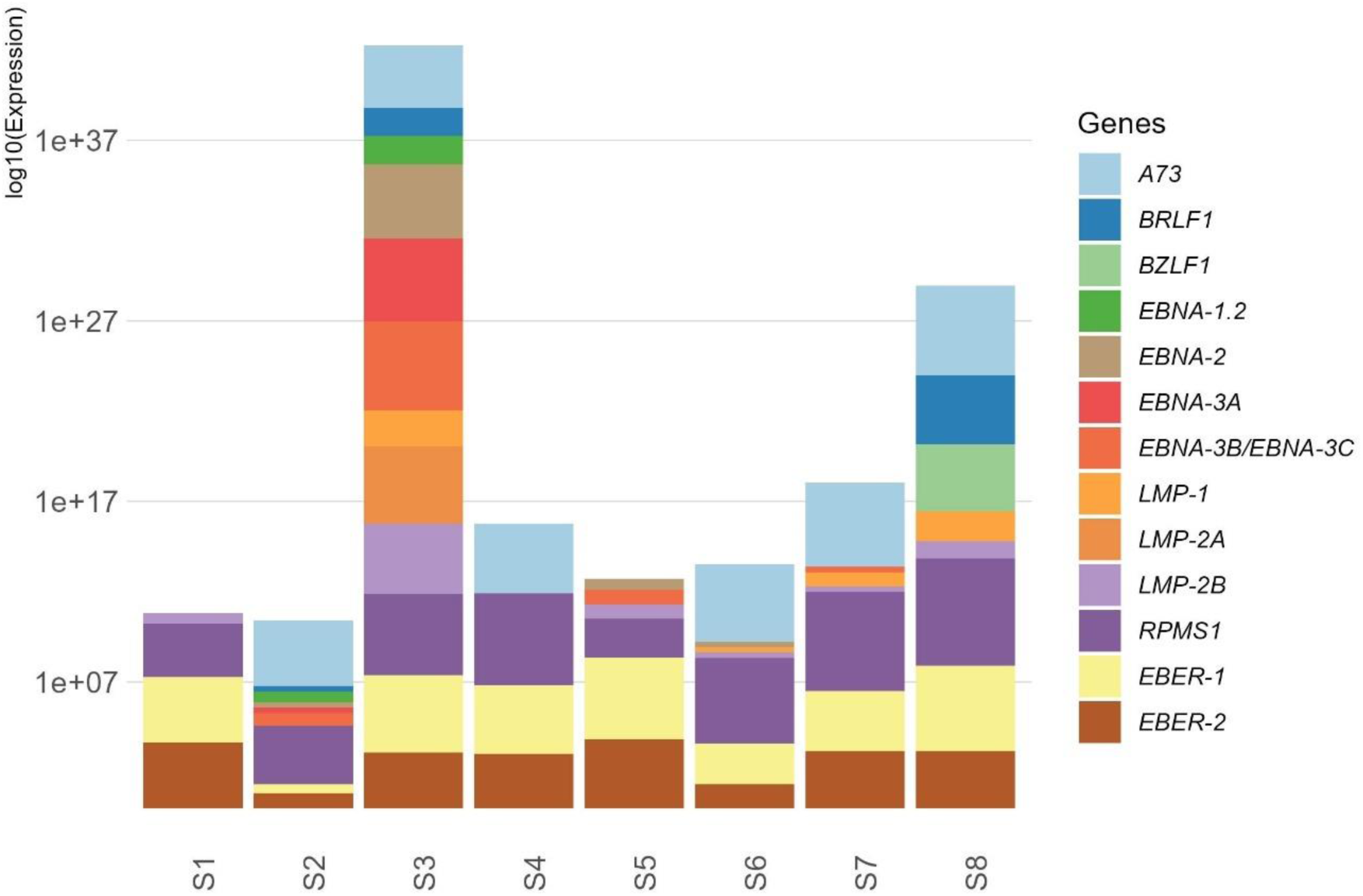
Expression of EBV viral genes by sample.

### Correlation of DE human gene expression with viral genes

After analyzing the expression of viral genes in positive samples, we correlated these genes with the previously analyzed DE human genes (**Fig 7**). Positive correlations between genes are shown in blue, while negative correlations are shown in red. We identified the formation of two main clusters: the first with a negative correlation between viral genes and DE genes and the second with a positive correlation. The first cluster comprises the following human genes: *PFKFB2, CLEC7A, GM2A, PARP14, CTSC, CXCL9, EPSTI1, PRPS2, GBP1,* and *TNFSF13B.* These genes are negatively correlated with viral genes from the *EBNA* family (*EBNA-1.2, EBNA-2, EBNA-3A, EBNA-3B/EBNA-3C)* and *LMP-2A*.

**Fig 7.**
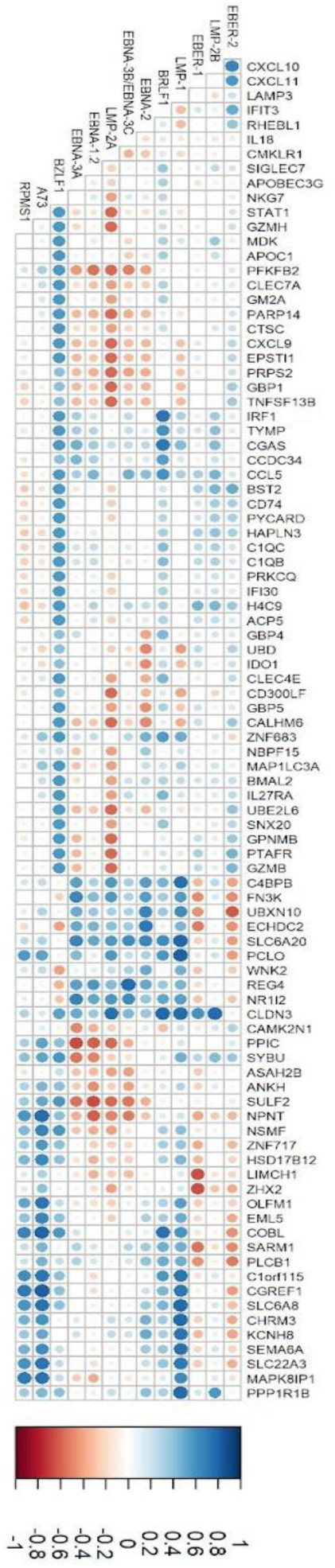
Correlation between DE genes with AUC > 0.85 and viral genes expressed in positive samples.

The second cluster includes the human genes: *C4BPB, FN3K, UBXN10, ECHDC2, SLC6A20, PCLO, WNK2, REG4, NR1I2,* and *CLDN3*. These genes are positively correlated with viral genes from the *EBNA* family *(EBNA-1.2, EBNA-2, EBNA-3A, EBNA-3B/EBNA-3C),* two genes from the *LMP* family *(LMP-1A* and *LMP-2A)*, and one gene from the lytic phase *(BRLF1*).

## Discussion

The identification of EBV presence in GC samples holds significant clinical relevance. Besides specific molecular and histological characteristics, patients diagnosed with this viral infection require different therapeutic approaches. In this context, we aim to evaluate a molecular classification method for the EBVaGC subtype using RNA-Seq data, examine differential gene expression between EBVaGC and EBVnGC patients, and investigate key viral gene expressions in virus-positive samples.

According to Murphy et al. [17], the prevalence rate of EBV in gastric tumors is approximately 9%, making the molecular classification of EBVaGC extremely important for a comprehensive understanding of the interaction between the virus and human gene expression. This can provide valuable information for the development of targeted therapeutic strategies and deepen our knowledge of the molecular bases of EBV-associated diseases.

Several studies have investigated the presence of the Epstein-Barr virus in gastric cancer samples. However, the molecular classification methods and approaches to viral gene identification used in these studies differ from those employed in our research. For instance, Camargo et al. [18] and the TCGA [5] utilized whole genome sequencing, whole exome sequencing, and analyses of mRNA and miRNA, supplemented by validation through immunohistochemistry (IHC) and in situ hybridization (ISH) tests. In contrast, our study focuses on RNA-Seq data analysis, complemented by ISH testing, to validate our findings.

Our results show that of the 76 samples analyzed, 8 were positive for the virus through the RNA-Seq method, corresponding to approximately 10.52% of the gastric tumors studied, corroborating the literature on EBV prevalence in gastric tumors[18,19]. A differential gene expression analysis of human genes between EBV-positive and EBV-negative samples was conducted. Approximately 834 genes were considered DE, and 92 of them showed an AUC > 0.85. Of these 92, we selected genes with the highest AUC values to infer their relationship with carcinogenesis or EBV and to determine if they could be considered biomarkers of EBVaGC.

The *GBP5* gene is crucial for activating the NLRP3 inflammasome, which is essential for innate immunity by signaling infections or tissue damage[20]. Comparative studies have shown its downregulation in EBVaGC samples compared to EBVnGC samples[21], and our findings confirm this, with an AUC greater than 0.95. Similarly, the *GBP4* gene, associated with increased inflammation in the tumor microenvironment—a key feature of EBVaGC—was also upregulated[6]. *CD274,* a gene that inhibits antitumor immunity by interacting with PD-1 on T cells, was also highly expressed, suggesting its role in immune evasion and its potential as a target for immunotherapy in EBV-associated cancers[22].

The *TYMP* gene also showed positive regulation in EBVaGC samples, with an AUC > 0.94. Although there is no specific literature linking it to EBV or the EBVaGC subtype, it is associated with gastric cancer (GC), where *TYMP* expression may influence tumor mutation load and correlate positively with the presence of CD8+ T cells[23]. This suggests that *TYMP* hyperexpression in EBV-positive samples might enhance CD8+ T cell infiltration, a characteristic frequently observed in EBVaGC samples compared to EBVnGC[24]. Further investigation is needed to understand *TYMP’s* role in the EBVaGC tumor microenvironment.

Although the gene *HAPLN3* was highly expressed in our positive samples, no associations were found with EBV or EBVaGC. However, this gene has been linked to other cancers, such as hypermethylation of its promoter in prostate cancer[25] and its role in cell proliferation, migration, and invasion in renal cancer[26]. In our research, some genes showed no direct or indirect associations with EBVaGC, EBV, or GC. Among them, the *GM2A* gene, despite its hyperexpression in positive samples, is primarily associated with degenerative diseases and not cancer. Conversely, *CGREF1*, known for its role in colorectal cancer prognosis and its involvement in regulating cell growth and proliferation, showed downregulation in our EBVaGC samples[27,28].

The *BST2* gene, as observed in this study, has shown positive regulation in GC samples and is associated with promoting epithelial-mesenchymal transition (EMT) and metastasis in GC when induced by *HOXD9*[29]. However, no connection was found between EBVaGC and EBV. Similarly, *CTSC* has been linked to cell migration, invasion, and EMT in glioma cells[30], as well as promoting breast cancer metastasis to the lung through neutrophil regulation[31]. However, no literature describes its role in EBVaGC or GC. Furthermore, the *GZMB* gene is known to influence GC cellular biology by promoting cell growth, migration, and EMT[32]. In our study, *GZMB* was upregulated in EBVaGC samples, suggesting an active immune response against tumor cells, as *GZMB* is a protease that induces apoptosis in target cells when released by cytotoxic lymphocytes or natural killer (NK) cells[33].

*SLC22A3* was downregulated in EBVaGC samples, contrasting with its overexpression in colorectal cancer cell lines, where it is linked to enhanced cell migration and invasion[34]. While the gene has been associated with various cancers, no connection with EBV or gastric cancer (GC) has been reported. Similarly, *HSD17B12*, also negatively regulated in positive samples, lacks documented associations with EBVaGC or GC. A study suggested its involvement in pathways linking inflammation to cancer, but further research is required to substantiate this[35]. The *SLC6A8* gene was also downregulated in our samples. Although no direct links to EBVaGC or GC have been identified, its role as a creatine transporter could be crucial in the energy metabolism of cancer cells, potentially affecting tumor progression[36].

*WNK2* has been characterized as a tumor suppressor, with epigenetic silencing reported in various cancers, including hepatocellular carcinoma[37]. In our study, *WNK2* was downregulated, suggesting that its reduced expression might promote tumor progression, supported by evidence that methylation by *LINC00858* restricts apoptosis and enhances invasion and migration in GC cells[38]. *CAMK2N1* was also found to be downregulated in EBVaGC-positive samples. Given that EBVaGC is associated with a favorable prognosis, its low expression could serve as a prognostic biomarker, as high expression of *CAMK2N1*, mediated by non-coding RNAs, is linked to poor prognosis and increased cell migration in GC[39]. Lastly, *NSMF*, though downregulated in our study, has not been previously associated with EBVaGC or GC. However, the deregulation of *NSMF* is known to cause chromosomal instability.

The results discussed so far suggest a fundamental role of DE genes in tumor development and progression. To better understand the function of these genes in GC biology, a gene ontology analysis was conducted, revealing that most DE genes are associated with biological processes related to the immune system and organismal defense.

The positive regulation of cytokine production is an important process for the immune system. It involves the participation of various genes that encode proteins necessary for cell signaling and the inflammatory response. Furthermore, cytokines can play significant roles in cancer development and in controlling distinct stages of the disease through interleukins, colony-stimulating factors, transforming growth factors, interferons, and chemokines[40]. An important and overexpressed cytokine among our DE genes is *IFNG*. This positive regulation can lead to an increase in the production of other cytokines, as it is one of the genes involved in the regulation of cytokine production and the positive regulation of the defense response. Its function as a gene may contribute to the innate immune response by reprogramming macrophages to the pro-inflammatory M1 phenotype and promoting T cells to produce additional cytokines that can enhance the immune response against tumor cells[41].

We know that the immune response includes both positive and negative regulations, which are important for inflammatory control and the prevention of autoimmune diseases. In this context, the gene ontology analysis showed that there are DE genes involved in the process of negative regulation of the immune system, among which we can highlight *PDCD1LG2/PD-L2*. *PD-L2* is a ligand for the *PD-1* receptor, a protein expressed on T cells that is associated with immune exhaustion, one of the mechanisms of negative immune regulation. Immune exhaustion can be characterized by the loss of T cell functions, potentially leading to apoptosis of these cells[42].

Another biological process highlighted in gene ontology is the regulation of T-cell activation, which is an intricate process involving transcription factors, various signaling cues, and regulatory mechanisms[43]. In the context of GC, regulatory T cells (Tregs) play a crucial role in the pathogenesis and maintenance of the central balance between immune suppression and autoimmunity. Additionally, Tregs are important in regulating T cell proliferation, as they can limit the expression of helper T cells and cytotoxic CD8+ T cells, promoting suppressive effects[44].

The biological processes of leukocyte-mediated immunity, regulation of leukocyte proliferation, and leukocyte proliferation are essential for both the adaptive and innate immune systems. Leukocytes are divided by types and functions, including lymphocytes, specifically T and B cells. Leukocyte-mediated immunity can occur through mechanisms such as antigen recognition, production, and activation of cytokines and chemokines, which are fundamental for communication between immune cells and for the recruitment of leukocytes to sites of inflammation or tumors[45]. The presence of ontology terms related to the adaptive immune response in our samples indicates that the presence of EBV in the tumor microenvironment may induce a strong antiviral immune response. This response involves the reactivation of defense cells that can combat both the virus and tumor cells. Moreover, it has been recorded that EBV employs various strategies to evade both innate and adaptive immune responses. However, in certain contexts, especially in latent infections, the immune system may be capable of detecting and responding to the virus by mobilizing CD8+ T cells and other adaptive immune responses to control the infection and potentially attack virus-associated tumor cells[46].

EBV encodes several proteins, and the coding patterns associated with several types of malignancies are variable. Moreover, the latent infection of the virus can be divided into four phases, with each phase expressing different viral genes. EBVaGC has been correlated with only one latency of the infection, latency I; however, recent studies also link it to latency II due to the expression of genes from this phase[8]. In this context, our analyses aimed to evaluate which viral genes were most expressed in our positive samples and what influence these genes may have on gastric cancer oncogenesis. According to our findings, *EBER-1* and *EBER-2* were expressed in 100% of the positive samples, confirming what has been described in the literature. These viral products are small non-coding RNAs expressed in all phases of latent infection and are used as diagnostic markers for detecting EBV[47]. We can directly associate the expression of *EBERs* with an increased immune response, as Kondo et al.[48] state that the loss of *EBER* expression is associated with decreased expression of *PD-L1* and CD8+ T lymphocytes. This suggests that by downregulating or reducing viral antigens, the tumor may escape immune surveillance, thereby facilitating tumor progression[48].

*BART* transcripts are described as a combination of microRNAs (miRNAs) and long non-coding RNAs (lncRNAs) expressed during the latent phase of infection. Their functions are associated with immune evasion and the promotion of carcinogenesis[6]. Among the results obtained, transcripts such as *RPMS1* and *A73* were identified. A previous study suggests that *RPMS1* is expressed only under specific conditions and may play a role in tumorigenesis, indicating that this transcript may be present during precancerous stages but not in malignant cells[49]. Similarly, *A73* demonstrated in vitro biochemical activities that may be relevant to carcinogenesis[50]. Our results show the manifestation of these viral products, *RPMS1* and *A73*, were detected in 100% and 75% of the samples, respectively, suggesting that these genes may be involved not only in the precancerous stage but also in contributing to the maintenance of carcinogenesis.

Additionally, we found genes from the lytic phase of infection expressed, known as immediate-early genes (IE), such as *BRLF1* and *BZLF1*, manifested in 25% and 12.5% of the samples, respectively. It has been demonstrated that *BZLF1* can alter the expression of several host genes and act by suppressing inflammatory responses to facilitate viral replication. In contrast, *BRLF1* can regulate both viral DNA replication and mRNA transcription[51]. Although there are no studies associating these genes with EBVaGC, understanding these genes and their functions may provide essential information for developing therapeutic strategies against EBV infection.

We also found the expression of viral genes from the *LMP* family, including *LMP-1, LMP-2A,* and *LMP-2B*. The manifestation of these genes in the samples can be explained by the fact that *LMPs* have oncogenic properties and the potential to transform epithelial cells by activating signaling pathways and modifying the expression of oncogenes and tumor suppressors[52]. In our results, *LMP-1* is expressed in 50% of the positive samples, although at low levels. This is consistent with the literature, which argues that despite being an important oncogene, *LMP-1* expression in gastric cancer is low or nonexistent due to the latency phase in which it is expressed[53]. Additionally, the low expression of this gene could be explained by a study suggesting that it may have been silenced through promoter methylation[52].

On the other hand, *LMP-2A*, found in 53.8% of EBVaGC cases described in the literature, is expressed in only 12.5% of our samples. One of the actions of *LMP-2A* is the activation of *STAT3* protein phosphorylation, which can bind to the enzyme DNA methyltransferase 1 (DNMT1). When this binding is positively regulated, it results in the silencing of tumor suppressor genes through CpG island methylation in their promoter region. These epigenetic changes, particularly DNA hypermethylation in the host genome, represent one of the main mechanisms involved in EBVaGC carcinogenesis[19,54]. In a systematic review of viral gene expression, Ribeiro et al.[55] reported that *LMP-2B* manifestation occurs in only 10% of cases. Our results show *LMP-2B* expression in 75% of our samples. With the low expression of this gene in EBVaGC cases, there is also a lack of studies on its role in gastric cancer development. Some studies describe that *LMP-2B* has the potential to regulate *LMP-2A* function, preventing potential EBV lysis. At the same time, high levels of *LMP-2B* expression may also accelerate the transition from latent to proliferative EBV infection[56].

Finally, we also observed the expression of genes from the *EBNA* family in our results. Among these, we found *EBNA 1.2*, which can be considered a variant of *EBNA1*, with low expression in 25% of our samples. This gene is considered essential for viral DNA replication and episome maintenance during cellular replication. Additionally, it regulates the expression of other EBV genes, controlling the transcription of viral genes that are important for latency[57]. The Epstein-Barr virus (EBV) is the first herpesvirus identified to be associated with human cancers known to infect the majority of the world population. EBV-associated malignancies are associated with a latent form of infection, and several of the EBV-encoded latent proteins are known to mediate cellular transformation. These include six nuclear antigens and three latent membrane proteins (LMPs). In lymphoid and epithelial tumors, viral latent gene expressions have distinct pattern. In both primary and metastatic tumors, the constant expression of latent membrane protein 2A (LMP2A) at the RNA level suggests that this protein is the key player in the EBV-associated tumorigenesis. While LMP2A contributing to the malignant transformation possibly by cooperating with the aberrant host genome. This can be done in part by dysregulating signaling pathways at multiple points, notably in the cell cycle and apoptotic pathways.

Recent studies also have confirmed that LMP1 and LMP2 contribute to carcinoma progression and that this may reflect the combined effects of these proteins on activation of multiple signaling pathways. This review article aims to investigate the aforementioned EBV-encoded proteins that reveal established roles in tumor formation, with a greater emphasis on the oncogenic LMPs (LMP1 and LMP2A) and their roles in dysregulating signaling pathways. It also aims to provide a quick look on the six members of the EBV nuclear antigens and their roles in dysregulating apoptosis[57]. The *EBNA-2* gene was found to be expressed in 50% of our results. The expression of this gene occurs soon after B cell infection by the virus. In a study published in 2023 by Zhang et al.[58], there is a suggestion that the expression of this gene, along with *LMP1* expression, may alter the transformation of primary B cells into lymphoblastoid cell lines[58].

There is no description in the literature of the presence of *EBNA-2* in the EBVaGC subtype, as it is typically expressed in latency phase 3. In contrast, the EBVaGC subtype expresses genes from phases 1 or 2. Other genes not reported as described in EBVaGC are those belonging to *EBNA-3*. However, in our samples, we found *EBNA-3A* expressed in 25%, with a high expression level in only one sample, and *EBNA-3B/EBNA-3C* expression in 50%. These genes function as transcription regulators of the virus and host cell, interacting with transcription factors to either transactivate or repress gene expression. Their manifestation is related to latency phase 3[57].

In summary, our findings confirm the expression of several EBV viral genes in positive samples, aligning with previous literature descriptions while also revealing a new perspective on the presence and potential impact of these genes in cancer cells. The expression of *EBER-1* and *EBER-2*, as well as the presence of *BART* transcripts, *RPMS1*, and *A73*, suggest a continuous contribution to the maintenance of carcinogenesis. The identification of lytic phase genes, such as *BRLF1* and *BZLF1*, in some samples, along with the manifestation of *LMPs* and *EBNAs*, highlights the complexity of EBV latent infection and the diversity of its gene expression patterns in gastric cancer. Given this, we sought to correlate viral genes and their expressions with differentially expressed human genes. We did not find studies directly describing the correlation of DE genes with viral genes, so it became necessary to understand which mechanisms the genes present in the identified clusters are associated with.

Among the genes present in cluster 1, we found some that are associated with important biological pathways, such as immunity, inflammation, and antigen presentation. In the immunity and inflammation pathway, the related genes are *CLEC7A, CXCL9,* and *TNFSF13B*[59–61], which are involved in the activation and survival of immune cells. In this regard, we can suggest that the expression of viral genes such as *EBNA-2* and *LMP-2A* may negatively influence the regulation of these genes, as the expression of these viral products can alter B cell transformation[58]. In addition to the mentioned genes, we can also suggest that the *PFKFB2* gene, involved in cellular metabolism regulation, may be influenced by the activity of transcription factors activated by viral genes such as *EBNA-2*[58,62].

In cluster 2, we found at least 10 DE genes that are positively correlated with viral genes; however, there are no descriptions explaining these correlations. We can infer about some genes, such as *PCLO*, which is associated with synapses and can influence cellular communication, as well as activate immune cells, which is crucial in the response to viral infections[63]. Additionally, the *REG4* gene is related to the inflammatory response and may interact with inflammatory processes induced by viral genes[64]. The WNK2 gene, previously suggested as a tumor suppressor, may be positively regulated by *LMP-2A* and its epigenetic functions of *STAT3* activation, as described in this study[37,54].

The expression of *BRLF1*, a gene active during the lytic phase of the viral cycle, may also modulate mRNA transcription[51]. Although the viral gene *BZLF1*, which is similarly involved in the lytic phase, is absent from any of the clusters described above, it demonstrates a positive correlation with the majority of differentially expressed human genes. This correlation can be explained by the function of *BZLF1*, as described in the present work, where the viral product can modify the expression of several host genes and suppress inflammatory responses, thus facilitating viral replication[51]. Although the human genes mentioned do not have a widely studied direct relationship with viral genes, their functions suggest that they may be modulated by viral proteins, affecting host response and disease progression. In this regard, further research is needed on these correlations since the interaction between human genes and EBV viral genes is complex and multifaceted.

In summary, this study conducted the molecular classification of EBV in gastric cancer using RNA-Seq data, validated by ISH assays, identifying eight positive samples. Furthermore, analysis of the gene expression in patients revealed 834 DE genes, 93 of which had an AUC > 0.85. These genes were associated with growth factor signaling, cell cycle regulation, metastasis, and biological processes linked to immune response and defense mechanisms. The viral gene expression of EBV was also analyzed in the positive samples, where we observed not only the expression of latency phase genes typically associated with EBVaGC but also the expression of viral replication phase genes. Correlating these viral expressions with the DE genes revealed modulation by viral genes that may impact the host immune response and disease progression. Although further investigations are needed better to understand the role of EBV in gastric carcinoma, our study presents an efficient strategy for the molecular classification of the EBVaGC subtype based on NGS and sheds light on the effects of EBV on human gene expression.

## Materials and methods

### Ethics statement and sample collection

This study was approved by the Ethics and Research Committee of João de Barros Barreto University Hospital (approval number: 47580121.9.0000.5634) and was conducted in accordance with the principles outlined in the Declaration of Helsinki. Participant recruitment and sample collection were carried out between July 2, 2022, and July 6, 2023. A total of 76 tumor tissue samples were obtained from patients diagnosed with GC who underwent surgical resection. Before enrollment, all participants were provided with detailed information about the study’s objectives, potential benefits, risks, and possible harms, ensuring a thorough understanding of the research. Written informed consent was voluntarily obtained from all participants prior to their inclusion in the study.

### Total RNA extraction

Approximately 30mg of tumor tissue from each sample was macerated for RNA extraction using the TRIzol Reagent method (Thermo Fisher Scientific), following the manufacturer’s instructions. The total RNA was then assessed for integrity and concentration (ng/μL) using the Qubit 4 Fluorometer (Thermo Fisher Scientific). The established criteria for the RNA Integrity Number (RIN) were ≥ 5.

### Library construction and RNA sequencing

Library construction was performed using the TruSeq Stranded Total RNA Library Prep Kit with Ribo-Zero Gold (Illumina, US) according to the manufacturer’s instructions, utilizing approximately 1 μg of total RNA per sample in a total volume of 11 μL. Upon completion of the libraries, the samples were assessed for the integrity of the generated DNA fragments using the 2200 TapeStation System (Agilent Technologies AG, Basel, Switzerland). Sequencing was performed in paired-end mode on the NextSeq® 500 platform (Illumina®, US), using the NextSeq® 500 MID Output V2 kit - 150 cycles (Illumina®) according to the manufacturer’s instructions. After sequencing, the obtained RNA-Seq reads were subjected to quality control. Initially, the quality of the reads was assessed using FastQC. Subsequently, adapters and low-quality reads were removed using Trimmomatic v0.39, with a quality value (QV) threshold set above 15.

### Molecular classification of EBVaGC

For the molecular classification of EBVaGC, the reads were aligned using the Kraken2 tool, utilizing the Reference Viral Database (RVDB) [15] as the index. The RVDB is a database with high specificity for identifying viral sequences in genomic and transcriptomic data.

### *In Situ* Hybridization

A tissue microarray library was constructed from gastrectomies of FFPE tissue archives. Representative tumor areas were selected by H&E review and applied to a new-generation tissue matrix (ngTMA Grand Master, 3DHistech) in duplicate and triplicate. The sections were subjected to in situ hybridization (ISH) reactions on the Dako automated system, detecting EBER1 RNA-1 (Y5200, DAKO, Carpinteria). Nuclear stain interpretations of the tests attributed positive staining to each case that presented a blue reaction.

### Alignment of samples with the human genome

Human transcript reads were characterized through alignment and quantification using Salmon v1.5.2, with the coding transcripts in hg v38 (www.ensembl.org) used as the reference index and GENCODE v.42 (www.gencodegenes.org) used for annotation. The reads were imported from Salmon into Rstudio using the Tximport v3.14.0 package.

### Differential gene expression and functional enrichment analysis of human genes

After classification and alignment, a differential gene expression analysis was performed between EBVaGC tumor samples and EBVnCG (non-EBV-associated CG) using the DESeq2 package. Genes considered differentially expressed (DE) were those whose expression difference met the criteria: i) | Log2(Fold-Change) | > 1; and ii) adjusted p-value < 0.05. The results of the differential gene expression were visualized using the ggplot2 v3.5.1 and complexHeatmap v2.14.0 packages. DE genes with AUC > 0.85 were enriched using Gene Ontology for terms related to biological processes, aiming to understand their biological and functional relevance better. The Cluster Profile v4.8.3 package was used for this analysis.

### Viral gene expression analysis

To assess the expression of EBV viral genes, samples identified as positive according to the RNA-Seq method were selected and processed using the *nf-core/RNAseq* v3.14 pipeline[16]. The EBV reference genome GCF_002402265.1 and genomic annotation NC_007605.1, both available from the National Center for Biotechnology Information (NCBI) database, were used as indices.

### Correlation of human DE gene expression with viral genes

A Spearman correlation was performed to assess the correlation between patient gene expression and expressed viral genes. The corrplot package in RStudio was used to visualize the correlations graphically.

## Author contributions

**Conceptualization:** Valéria Cristiane Santos da Silva, Diego Pereira, Fabiano Cordeiro Moreira, Paulo Pimentel de Assumpção, Rommel Mario Rodriguez Burbano;

**Data curation:** Valéria Cristiane Santos da Silva, Diego Pereira, Daniel de Souza Avelar, Fabiano Cordeiro Moreira;

**Formal analysis:** Valéria Cristiane Santos da Silva, Diego Pereira, Ronald Matheus da Silva Mourão, Fabiano Cordeiro Moreira;

**Funding acquisition:** Ândrea Kely Campos Ribeiro dos Santos, Fabiano Cordeiro Moreira, Geraldo Ishak, Samia Demachki, Samir Mansour Casseb, Paulo Pimentel de Assumpção, Rommel Mario Rodriguez Burbano, Williams Fernandes Barra;

**Investigation:** Valéria Cristiane Santos da Silva, Diego Pereira, Daniel de Souza Avelar, Jéssica Manoelli Costa da Silva, Ronald Matheus da Silva Mourão, Rubem Ferreira da Silva, Davi Josué Marcon, Leandro Magalhães, Amanda Ferreira Vidal, Fabiano Cordeiro Moreira; Samir Mansour Casseb;

**Methodology:** Valéria Cristiane Santos da Silva, Diego Pereira, Daniel de Souza Avelar, Jéssica Manoelli Costa da Silva, Ronald Matheus da Silva Mourão, Rubem Ferreira da Silva, Davi Josué Marcon, Leandro Magalhães, Amanda Ferreira Vidal; Tatiane Neotti, Ana Karyssa Mendes Anaissi, Samia Demachki, Samir Mansour Casseb, Fabiano Cordeiro Moreira;

**Sample Acquisition:** Geraldo Ishak, Paulo Pimentel de Assumpção, Rommel Mario Rodriguez Burbano, Williams Fernandes Barra;

**Supervision:** Fabiano Cordeiro Moreira, Samir Mansour Casseb, Paulo Pimentel de Assumpção, Rommel Mario Rodriguez Burbano;

**Project Administration:** Fabiano Cordeiro Moreira, Samia Demachki, Samir Mansour Casseb, Paulo Pimentel de Assumpção, Rommel Mario Rodriguez Burbano;

**Validation:** Tatiane Neotti, Ana Karyssa Mendes Anaissi, Samia Demachki, Williams Fernandes Barra;

**Visualization:** Valéria Cristiane Santos da Silva, Diego Pereira;

**Writing – original draft:** Valéria Cristiane Santos da Silva, Diego Pereira, Fabiano Cordeiro Moreira;

**Writing – review & editing:** Valéria Cristiane Santos da Silva, Diego Pereira, Daniel de Souza Avelar, Jéssica Manoelli Costa da Silva, Ronald Matheus da Silva Mourão, Rubem Ferreira da Silva, Davi Josué Marcon, Leandro Magalhães, Amanda Ferreira Vidal; Tatiane Neotti, Ana Karyssa Mendes Anaissi, Williams Fernandes Barra, Samia Demachki; Samir Mansour Casseb, Andrea Kely Campos Ribeiro dos Santos, Paulo Pimentel de Assumpção, Fabiano Cordeiro Moreira, Rommel Mario Rodriguez Burbano.

## Acknowledgment

The authors would like to thank the Oncology Research Center, the Human and Medical Genetics Laboratory, and the Anatomical Pathology Laboratory at João de Barros Barreto University Hospital (HUJBB – UFPA) for their invaluable technical and laboratory support. Our gratitude also goes to the High-Performance Computing Center (CCAD) at the Federal University of Pará for access to the Apollo 2000 cluster, which was crucial for our analyses.

